# A simple age-structured, temperature-dependent population growth model of brown rice hopper (*Nilaparvata lugens* (Stål))

**DOI:** 10.1101/2025.01.15.633297

**Authors:** Xiaojing Wei, Jane Girly Balanza, Vu Thi Bich Ngoc, James Giles, Cornelis Swaans

## Abstract

Brown rice hopper (BRH), a key insect pest to rice, is known can change its longevity and fecundity due to rising temperature. Here we modeled the response of BRH population growth to temperature using an age-structured population growth model, based on reported responses of BRH development time, survival, and fecundity to temperature. We applied this model to predict historical level of BRH population growth in the Mekong River Delta (MRD) and compared the predictions with historically observed percentages of rice fields infested by BRH. The model was able to capture the seasonal variation in BRH infestation and the optimal temperature for BRH activities (app. 26.5°C). These results highlighted the importance of temperature in regulating BRH population growth. The model, with further improvements as discussed, could be used for projecting BRH activity under rising temperatures or predicting BRH outbreaks due to seasonal temperature anomalies.

## Introduction

Climate change alters the distribution of agricultural pests and their impact on crop yield Savary et al. 2006, Annu. Rev. Phytopathol.). For instance, rising temperature can influence the vital rates of insect pests. These temperature-induced changes, in turn, could lead to changes in the growth rate of pest populations and their impact on crop yield (Deustch et al. 2018, Sci.).

Brown rice hopper (hereafter BRH), *Nilaparvata lugens* (Stål), is a rice-monophagous, outbreak-prone (i.e., short generation, high reproduction rate and migratory capacity) sap feeder and pathogen transmitter that has become prominent since the Green Revolution due to excessive use of fertilizers and pesticides (Bottrell et al., 2012, J. Asia-Pac. Entomol.). Experimental studies have demonstrated that increasing temperature can alter individual BRH’s longevity, fertility, and body size (Horgan et al. 2020, PlosOne). Yet, few studies have synthesized these evidences to infer the impact of rising temperature on the growth and consumption of BRH at the population level.

Here we synthesized the reported experimental data and modeled the response of BRH population growth to temperature. This model, adapted from a generic model of (Deustch et al. 2018, Sci.), could be used to project the impact of BRH on crop yield under future climate scenarios. We applied this model to predict historical level of BRH population growth in the Mekong River Delta and compared the predictions with historically observed BRH infestation level.

## Methods

### Adapting the generic insect pest model of Deustch et al. for BRH

Deutsch et al. (2018, Sci.) modeled crop yield loss due to insect pests as the product between *per capita* pest consumption rate and pest population density. Their modeled consumption rate as proportional to the metabolic rate of the pest and the response of metabolic rate to temperature following the Kleiber’s rule. They modeled population growth using a logistic growth model, of which the intrinsic growth rate declining in proportion to the degree of deviation from the optimal temperature of the insect population. They parameterized the model using estimates of historical yield loss at the spatial resolution of UN sub-region.

We modified the model of Deustch et al. to model the responses of BRH growth to temperature. Instead of using a logistic growth model, we used an age-structured population growth model implemented using the DemogR package in R (Jones et al. 2007, J. Stat. Softw.).

Following Horgan et al. (2020, PLOS ONE), we modeled the response of longevity and lifetime fertility of female BRH adults to temperature as quadratic, concave functions. Also following Horgan et al., we modeled the age (i.e., day)-specific, accumulative survival and instantaneous fecundity of female BRH adults as linear, declining functions (i.e., both survival and fecundity decreased to zero at the end of the female lifespan). Combing these functions, we derived age-specific, survival and fecundity of female BRH adults as functions of temperature.

We modeled the responses of BRH nymph developmental time and accumulative survival over the entire nymph stage (l_x_, x= nymph developmental time) to temperature as quadratic, concave functions, pooling data from Horgan et al. (2020, PLOS ONE) and Rout & Jena (2012, ORYZA). Horgan et al. showed that the age-specific, instantaneous survival (S_x_) of BRH nymph was constant (and close to 100%) under 20°C, 25 °C, and 30 °C, but declined with age under 35 °C. In line with Horgan et al, Rout & Jena showed that the age-specific, accumulative survival (l_x_) of BRH nymph was 100% under 30 °C but decreased with age under higher temperature. They also showed that as temperature increased, the negative slope of l_x_ to age became steeper. Synthesizing these evidence, we modeled the age-specific survival of BRH nymph using the Weibull distribution model (Amarasekare et al. 2012, Funct. Ecol.). From 20 °C to 30 °C, we set the shape parameter of the Weibull distribution model to 1 (i.e., constant S_x_ across age). From 30 °C to 35 °C, we modeled the shape and scale parameter of the Weibull distribution model as linear functions of temperature. We determined the unknown parameters of the Weibull distribution models based on the accumulative survival over the entire nymph stage (l_x_, x= nymph developmental time). Finally, we assumed 100% survival from eggs to nymphs based on the experimental results of Horgan et al. (2020, PLOS ONE).

Combining the above-mentioned response functions of the vital rates of BRH female adults, nymphs, and eggs to temperature, we implemented R functions that generate lifetable and the corresponding Leslie matrix based on given temperature using the *life*.*table()* and the *leslie*.*matrix()* function in the demogR package.

### BRH population dynamics in Mekong River Delta

The Mekong River Delta (MRD), known as the “rice bowl” of Vietnam, contributes to 51% of the national rice production of Vietnam (Käkönen et al. 2008, AMBIO) – the third largest rice exporting country (Ha et al. 2015, JAE). In MRD, rice production usually has three cropping seasons: winter-spring (Dong-Xuan), app. from November to February; summer-autumn (He-Thu), app. from April to July; and monsoon (Mua), app. from August to November (Ferrer et al. 2022, EAP).

Otuka et al. (2014, Appl. Entomol. Zool.) found that in MRD, BRH goes through app. 12 generations per year (app. 4 generations per cropping season), and these generations were synchronized across the delta, suggesting migration played a minor role in shaping BRH population dynamics in MRD. Within a cropping season, BRH population density peaked in the middle of the season and crashed after the harvest. Within a year, peak BRH population density tended to be highest during the winter-spring or the summer-fall season.

### Preliminary model validation

We predicted BRH population growth during the winter-spring and the summer-autumn cropping season from 2019 to 2023 in six provinces in MRD (An Giang, Dong Thap, Kien Giang, Vinh Long, Soc Trang, Tien Giang) based on the mean temperature of the rice production area in the provinces over the projected periods. We obtained historical daily temperature from AgERA5 at a spatial resolution of 5 km. We obtained rice production map of the six provinces based on the 2020 MapSpam rice planting area data at a spatial resolution of 10km. We assumed that the population started at 10 reproducing female adults in the beginning of the cropping season and projected population size at the end of 70 days.

We obtained the area of BRH infested paddy during the winter-spring and the summer-fall season from 2019 to 2023 in each of the six provinces from the Plant Protection Department under the Ministry of Agriculture and Rural Development of Vietnam. We obtained the planted area of paddy during the same time periods of each of the provinces from the General Statistics Office under the Ministry of Planning and Investment. We calculated the percentage of the BRH infested paddy area by dividing the former by the latter.

## Results and discussion

### Response of simulated BRH population growth to temperature

Fig.1 showed the accumulative survival(l_x_), fecundity (f_x_), and reproduction number (R_x_) of the BRH model over a series of temperature from 25 to 30 °C. Nymph survival showed a quadratic, concave response to temperature, with an optimal temperature around 28°C (Fig.1a). Adult survival and fecundity also showed quadratic, concave response to temperature, with optimal temperature around 26°C(Fig.1b). Together, the responses of these vital rates caused R_x_ to show a quadratic, concave response to temperature, with an optimal temperature around 26.5 (Fig.1c).

**Fig 1.**
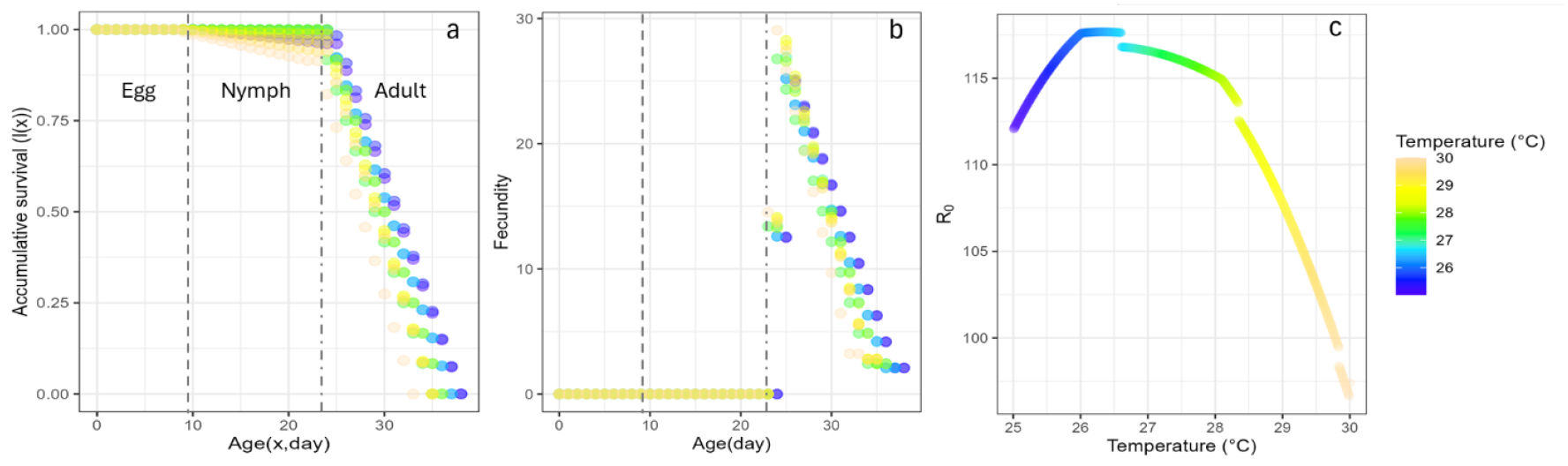
Survival (a), fecundity (b), and reproduction number (c) of the brown rice hopper model across a series of temperature.

Fig. 2 showed the projected BRH population growth over time. As expected for an age-structured population growth model with relatively long non-reproductive life history stages (app. two thirds of the entire lifespan), population growth showed distinct bursts, corresponding to the periods when new cohorts reached the reproductive age (Fig.2a). The responses of the stabilized population size after these bursts of reproduction to temperature followed the pattern of R_x_ (Fig. 2b-g).

**Fig 2.**
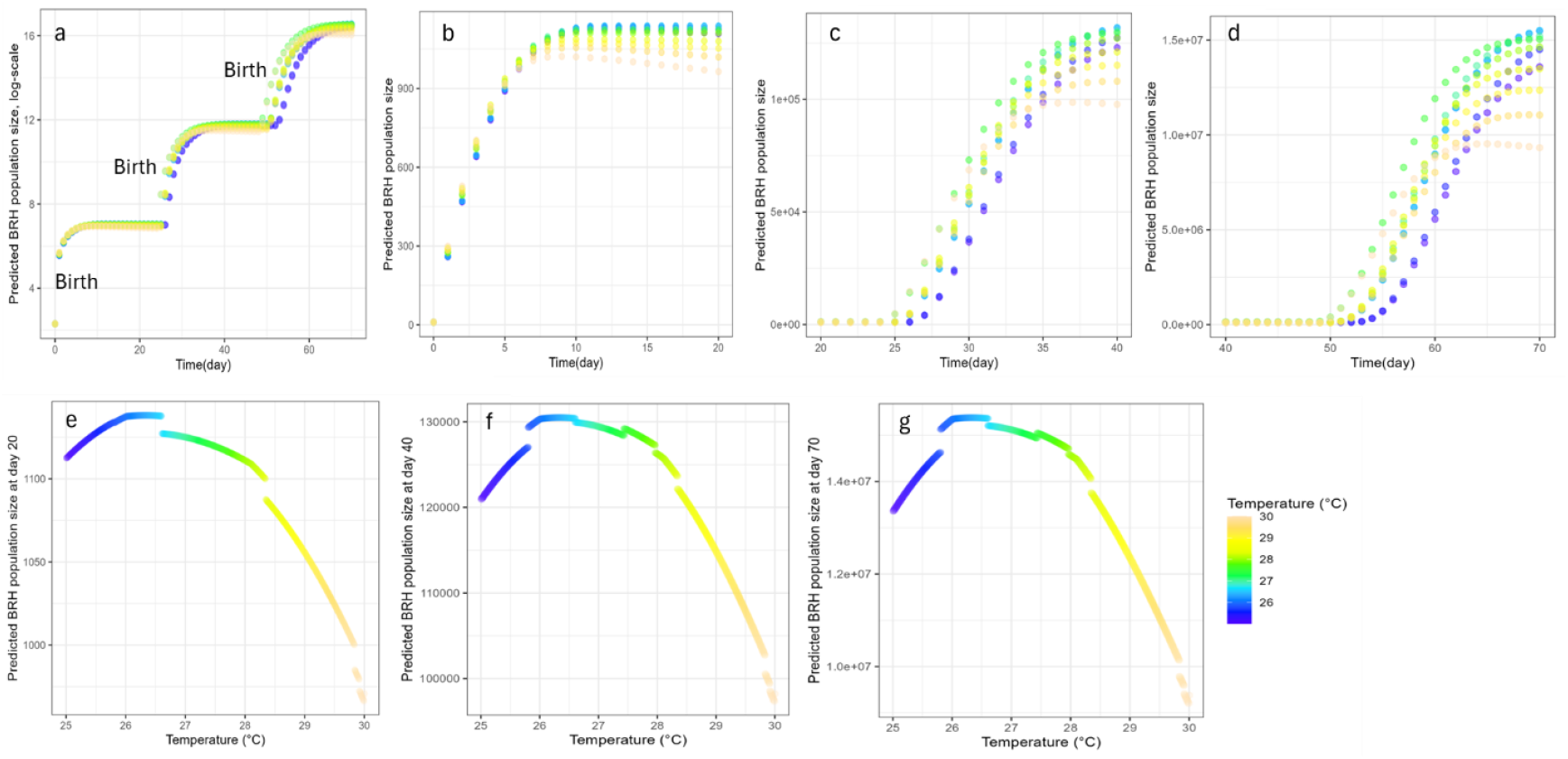
Projected brown rice hopper (BRH) population growth over time. Panel (a) showed BRH population size over time on a log scale. Panel (b)-(d) showed BRH population growth over 0 to 20 days, 20 to 40 days, and 40 to 70 days, respectively. Panel (e)-(g) showed BRH population size at day 20, 40, and 70.

### Preliminary model validation

Using a relatively small dataset (60 data points in total), namely, the percentage of BRH infested rice fields during the peak of the winter-spring and the summer-fall cropping season in six provinces in MRD from 2019 to 2023, we conducted a preliminary validation of our BRH model. The observed percentage of BRH infested rice paddies was significantly higher in the winter-spring than the summer-fall season (Fig.3a), showed a declining trend over year(Fig.4a), and was highest in An Giang province during the winter-spring season(Fig. 5a). We validated the model by examining if the projected BRH population size at day 70 showed similar patterns.

**Fig 3.**
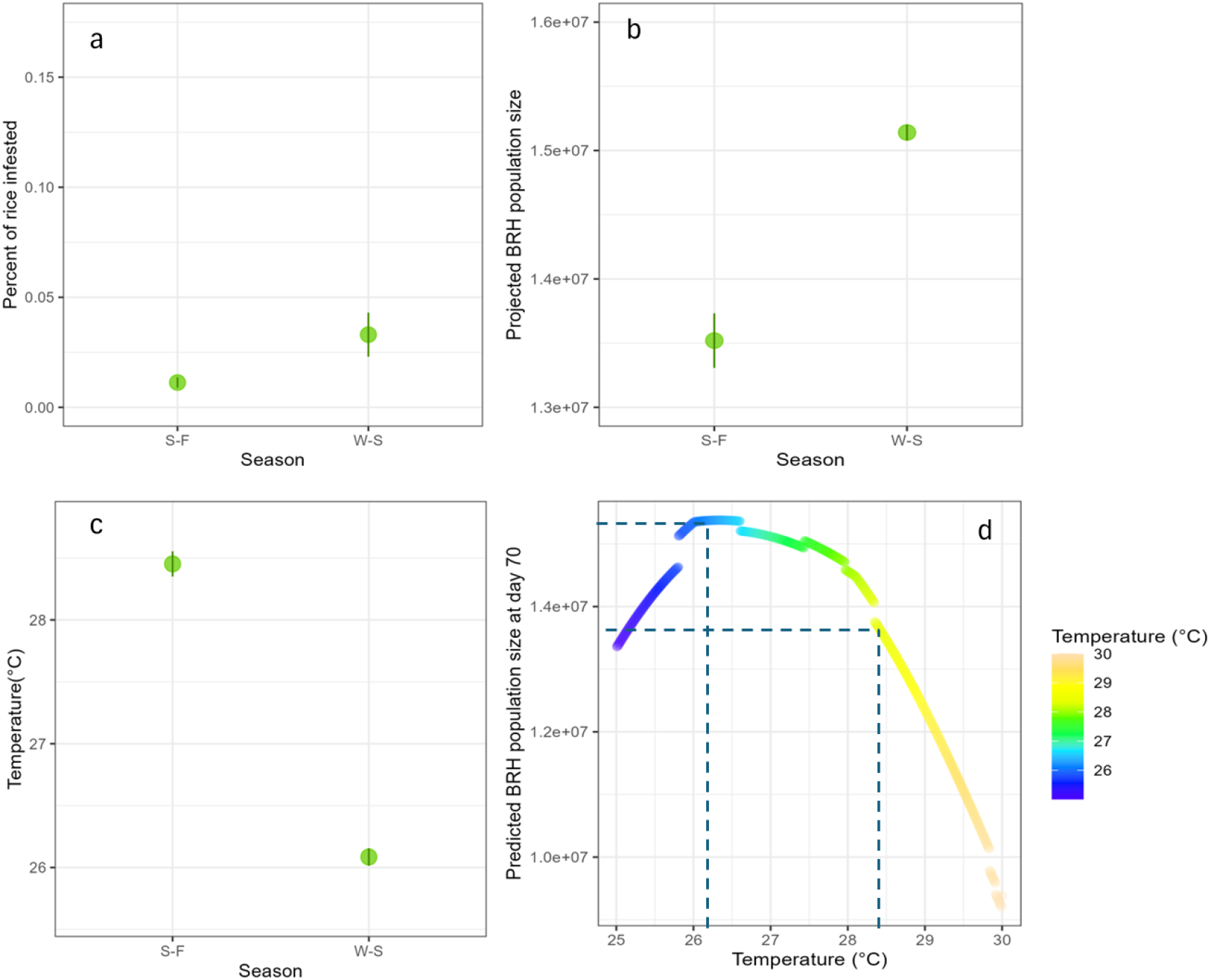
Seasonal variation in brown rice hopper (BRH) activity. Panel (a), (b), and (c) showed the percentage of rice fields infested by BRH, the projected BRH population size at day 70, and temperature in summer-fall (S-F) vs. winter-spring (W-S) seasons. The green dots and vertical lines indicate the mean and the standard errors of the means, respectively. Panel (d) showed the response of the projected BRH population size at day 70 to temperature.

**Fig 4.**
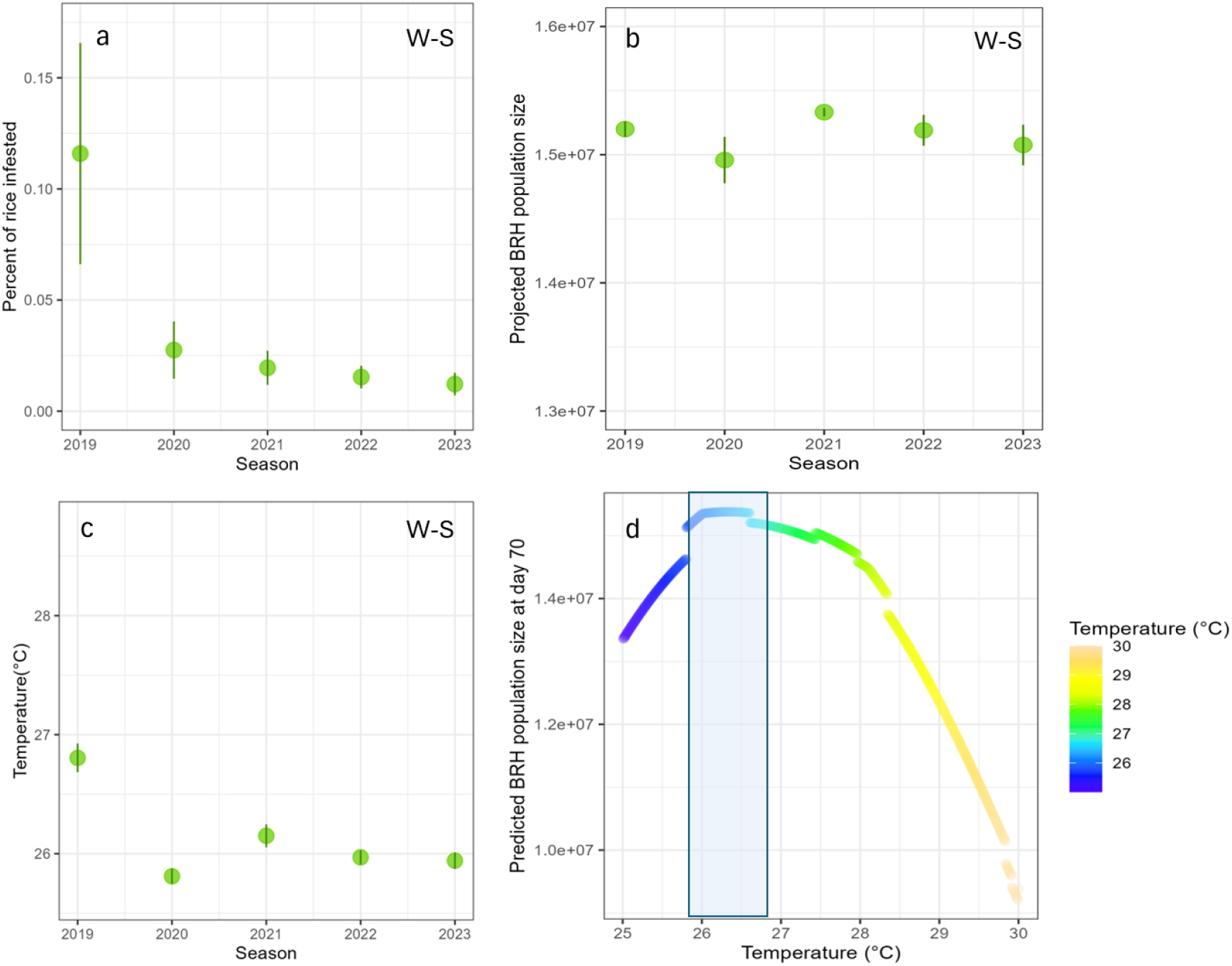
Among-year variation in brown rice hopper (BRH) activity. Panel (a), (b), and (c) showed the percentage of rice fields infested by BRH, the projected BRH population size at day 70, and temperature by year during the winter-spring (W-S) cropping seasons. The green dots and vertical lines indicate the mean and the standard errors of the means, respectively. Panel (d) showed the response of the projected BRH population size at day 70 to temperature.

**Fig 5.**
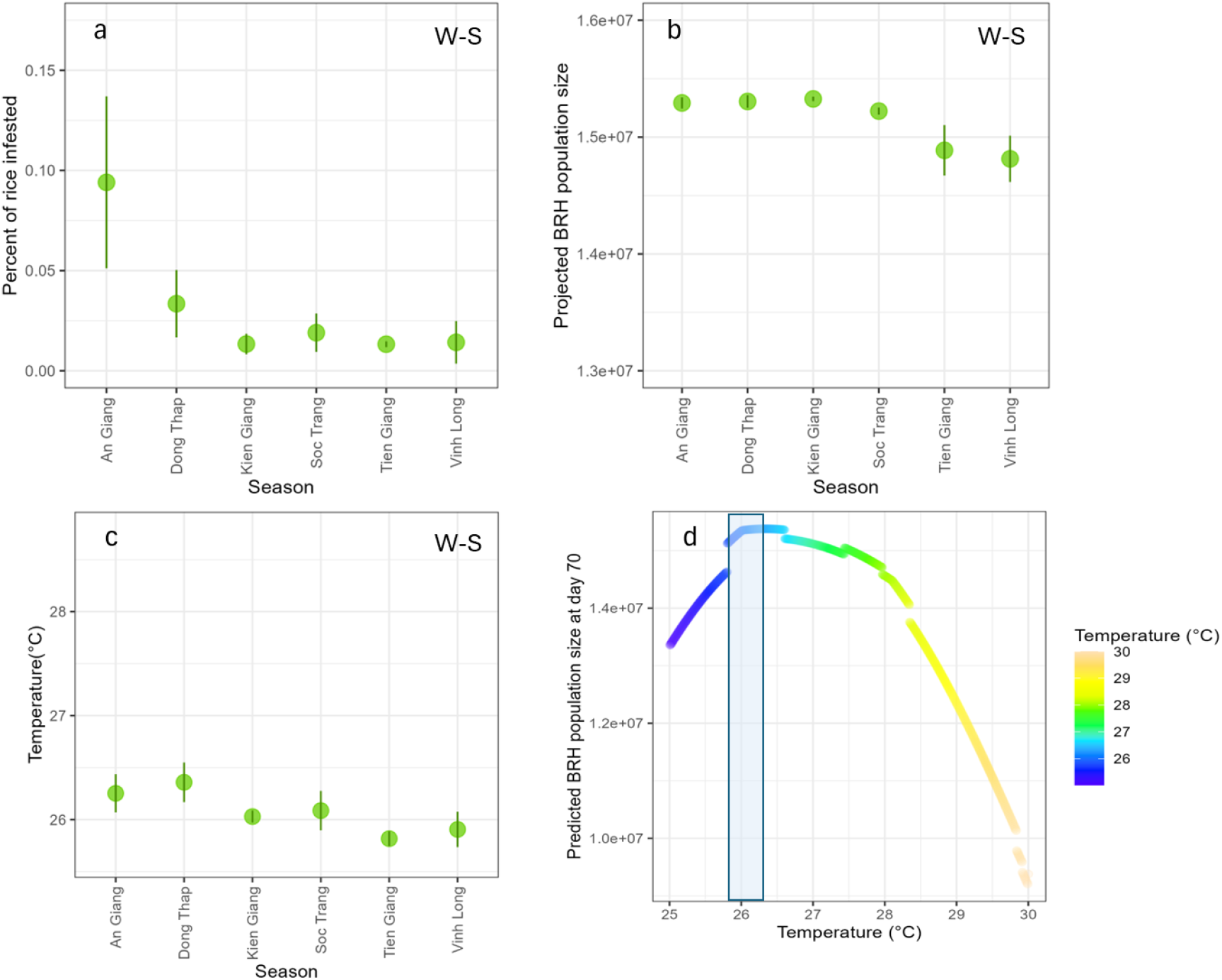
Among-province variation in brown rice hopper (BRH) activity. Panel (a), (b), and (c) showed the percentage of rice fields infested by BRH, the projected BRH population size at day 70, and temperature by province during the winter-spring (W-S) cropping seasons. The green dots and vertical lines indicate the mean and the standard errors of the means, respectively. Panel (d) showed the response of the projected BRH population size at day 70 to temperature.

In line with the observed data, the projected BRH population size at day 70 was significantly higher in the winter-spring than the summer-fall season (Fig.3b). However, unlike the observed data, it did not show a significant, declining trend over year (Fig. 4b), nor was it highest in the An Giang province (Fig. 5b). Three reasons might have contributed to these inconsistent validation results. First, the temperature difference between the winter-spring and the summer-fall season was relatively large (app. 2°C, Fig. 3c-d), while the temperature differences among year and province were smaller (app. 1 and 0.5°C, respectively; Fig.4c, Fig.5c) and were in a relatively flat range of the response of population growth to temperature (Fig.4d, Fig.5d). Second, the significant variations in the observed BRH infestation among year and province might have been driven by a single outlier – An Giang province in 2019 (Fig. 6a&c), as reflected by the relatively large error bars in Fig. 4a and Fig. 5a. Last, factors that not considered in our simple model might have contributed to the observed variations among year and province (see the *Future directions* section below).

**Fig 6.**
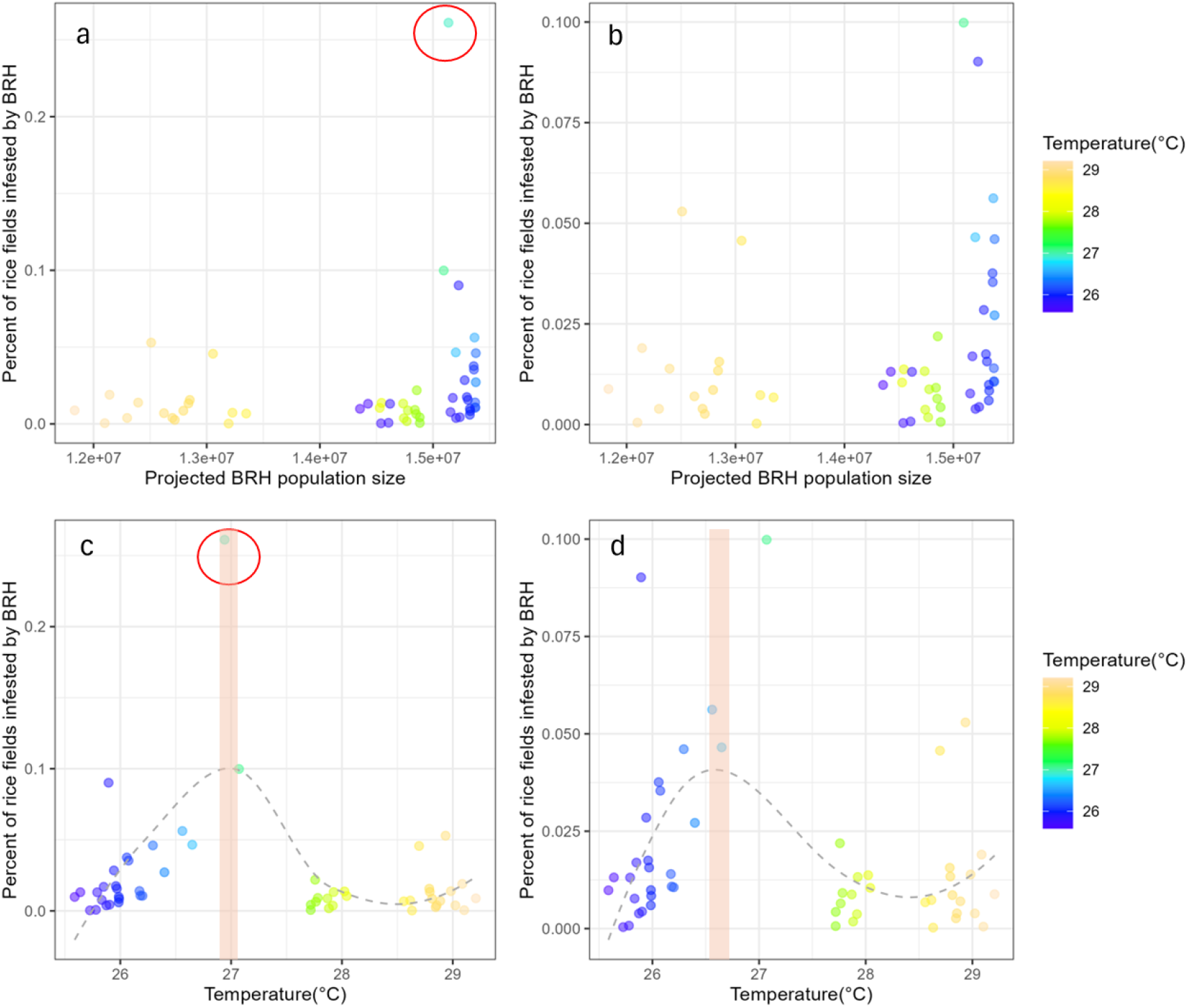
Comparison between the observed percent of rice fields infested vs. the projected BRH population size at day 70. Panel (a) and (b) showed the former plotted against the latter, with and without the outlier (highlighted in the red circle), respectively. Panel (c) and (d) showed the observed percentage of infested rice fields to temperature with and without the outlier, respectively. The dashed curves indicate the loess regression curves.

In line with the above-mentioned results, the Spearman’s correlation coefficient between the observed BRH infestation rate and the projected BRH population size at day 70 was 0.34 (P=0.01, Fig. 6a), indicating that our model captured some but not all of the variation in the observed data. Note that the rank correlation coefficient stayed 0.33 (P =0.02, Fig. 6b) after removing the outlier (An Giang, 2019, winter-spring season). Finally, the observed BRH infestation rate showed a quadratic concave response to temperature (Fig. 6c). After removing the outlier, the quadratic curve peaked around 26.5 °C (Fig. 6d), i.e., similar to the projected population growth and R_x_. However, the observed BRH infestation rate did not decline further beyond 28°C, while the projected population growth did.

In summary, our BRH model was able to capture the quadratic, concave shape of the observed BRH infestation rate to temperature and predict an optimal temperature that was relatively close (<0.25 °C) to the temperature at which the observed BRH infestation peaked. Furthermore, it captured the seasonal pattern in the observed BRH infestation. However, it did not discern more subtle variation among-year and among-province that occurred over smaller temperature variation (< 1°C).

### Future directions

Our model is intentionally simple and leaves out several factors important to BRH population growth, ranging from humidity to pest management practices. The fact that such a simple model could capture the optimal temperature for the observed BRH infestation and the pattern in its seasonal variation highlights the importance of temperature in regulating BRH population dynamics. It also suggests that our simple model holds the potential of further development into a more useful tool for projecting BRH infestation under future climate or for predicting the likelihood of BRH outbreak based on seasonal forecasts of unusually high temperature. In the spirit of making our model more useful, we propose the following future directions.

First, our preliminary validation could be greatly improved by more validation data distributed more evenly across the temperature gradient and with a longer time sequence (replicated across multiple locations). Such validation would allow us to better access under which condition the model perform well, which is necessary for designing proper user cases.

Additional data could also be used to calibrate the model, improving model accuracy or tailoring it to different user cases. For instance, our preliminary analysis suggested that BRH infestation varied among provinces. If such variation among provinces was consistent over a longer time sequence, then calibration could be used to tailor the model to different locations.

Second, currently our model is density-independent. A relatively simple way to incorporate density-dependency is to use the reported the time to which BRH populations peaked in the field (like those by Otuka et al.) in combination with the R_x_ of our current model to derive carrying capacity.

Last, in order to project BRH activities under climate in distant future, when temperature would rise substantially, it is important to understand why the model projection did not match the observed BRH infestation beyond 28 °C. Note that all the observed data beyond 28°C was collected during the summer-fall season. One possibility was that the BRH population dynamics differed between the season due to factors other than temperature, e.g., humidity.

## Acknowledgement

X. Wei thank K.-H. Kim and N. Castilla for insightful discussions that has substantially improved the work and T. T. Le for obtaining the BRH infestation data. This work is funded by the CGIAR initiative Asian Mega Deltas.

